# Representation of illusory shapes within the topographic areas of the posterior parietal cortex

**DOI:** 10.1101/2022.04.12.488016

**Authors:** Ana Arsenovic, Anja Ischebeck, Natalia Zaretskaya

## Abstract

The human visual system consists of multiple topographic maps that extend from the early visual cortex along the dorsal and ventral processing streams. Responses to illusory shapes within these maps have been demonstrated in the ventral stream areas, in particular the lateral occipital complex. Recently, the intraparietal sulcus of the dorsal stream has been linked to the processing of illusory shapes defined by motion. It therefore remains unclear whether the topographically organized parietal areas also respond to static illusory shapes, which would suggest their generic role in representing illusory content. Here we measured brain responses using fMRI while human participants observed flickering inducers around the fixation task. The inducers either formed an illusory diamond in the center, a triangle in the left or in the right hemifield, or were inverted such that no illusory figure was formed. We compared responses of parietal regions IPS0-IPS5 and SPL1 to each illusory figure with the non-illusory condition. To determine the role of attention in illusory shape responses we manipulated the difficulty of the fixation task. Our results show that all IPS areas responded to illusory shapes. The more posterior areas IPS0-IPS3 additionally displayed a preference towards the contralateral shapes, while the more anterior areas IPS4 and IPS5 showed response attenuation with increased task difficulty. We suggest that the IPS can represent illusory content irrespective of the perceptual mechanism that generated it. These responses may serve as a potential feedback signal that drives illusory shape responses in early and ventral visual areas.

**Significance statement:** The traditional view of the ventral visual pathway being solely responsible for representation of objects has recently been challenged by demonstrating illusory shape representation within the dorsal visual pathway with moving bistable stimuli. Our results provide evidence for the dorsal stream contribution to representing not only moving, but also static illusory shapes. Our results also show a functional subdivision along the topographic maps, with spatially specific shape responses in the more posterior, and attention-dependent responses in the more anterior areas. IPS areas of the dorsal stream should thus be considered in the theoretical accounts and neural models of how subjective content is generated in the brain.

## Introduction

One of the fundamental principles of brain organization is topography. Each sensory modality contains multiple topographic maps, which are considered to be central to information processing within their respective modalities (Kaas, 1997; Silver & Kastner, 2009). Functional magnetic resonance imaging of the visual system revealed that topographic organization is a property of many areas far beyond the early visual cortex (EVC). The topographic visual organization can be detected in mid-level visual areas located further laterally, ventrally and dorsally from the occipital lobe. Currently, more than 30 topographic maps have been identified across the visual processing hierarchy, extending from subcortical visually responsive areas (DeSimone et al., 2015; Schneider, 2004) and EVC (Engel, 1997; Sereno et al., 1995), up to parahippocampal areas along the ventral stream and the frontal eye fields along the dorsal stream (Wang et al., 2015). The exact functional role of these maps remains a topic of debate (Wandell et al., 2007).

Previous studies have shown that topographic maps in the EVC can represent not only simple sensory features (e.g., orientation, color, motion direction), but also more complex illusory content. For example, the EVC shows retinotopic responses not only to physical, but also to illusory contours (Larsson et al., 1999; Lee & Nguyen, 2001; Maertens et al., 2008; von der Heydt et al., 1984; Zhou et al., 2000). Furthermore, it shows response enhancement at the topographic representation of the illusory surface and suppression at the location of the inducers and the background (Grassi et al., 2017; Kok et al., 2016; Kok & de Lange, 2014). This complex response pattern in the EVC is thought to be a result of feedback modulation from higher-level areas, which also respond to illusory contours (Anken et al., 2018; Mendola et al., 1999; Murray et al., 2002).

According to the two-streams hypothesis, the perception of shapes, including illusory shapes, is a function typically attributed to the ventral stream (Goodale & Milner, 1992; Haxby et al., 1991). Indeed, multiple studies have demonstrated the involvement of the lateral occipital complex (LOC) in the perception of not only real, but also illusory shapes (Chen et al., 2020; de-Wit et al., 2009; Murray, 2004; Stanley & Rubin, 2003), making it a candidate area for providing the feedback signal to early visual areas (Fang et al., 2008; Wokke et al., 2013). However, the strict dichotomy between the ventral and the dorsal stream in the context of object perception is being continuously challenged (Freud et al., 2016). Recently, several studies reported activity of the posterior parietal cortex (PPC) and, specifically, the intraparietal sulcus (IPS) of the dorsal stream during perception of illusory shapes (Grassi et al., 2018; Zaretskaya et al., 2013; Zaretskaya & Bartels, 2015). These studies used bistable stimuli which can be interpreted either as an illusory global shape or as local non-illusory elements based on the same sensory input. Crucially, the illusory shapes investigated in these studies were defined by motion cues, which are expected to be processed along the dorsal stream. The PPC is known to contain multiple attention-defined topographic maps (Konen & Kastner, 2008; Schluppeck et al., 2005; Sereno et al., 2001; Silver et al., 2005; Swisher et al., 2007). It is therefore unclear whether the topographic areas of the PPC also respond to static illusory shapes and whether these responses depend on attention, which illusory shapes are known to capture (Nowack et al., 2021; Senkowski et al., 2005).

In this study, we investigated the illusory shape activity within the topographic maps of the PPC. Specifically, we were interested in whether parietal topographic areas respond to static illusory Kanizsa shapes and how these responses are modulated by top-down attention. We conducted an fMRI study where we measured brain responses of human participants while they viewed Kanizsa figures and performed a central fixation task. By independently manipulating the figure location and task difficulty, we were able to dissociate the shape-related from the attention-dependent parietal activity.

## Material and methods

### Participants

Thirty healthy volunteers (12 men), aged 19-31 years (*M* = 23.53 ± 2.92 *SD*), participated in the experiment. All but one of the participants were right-handed. All participants had normal or corrected-to-normal vision and did not take any psychotropic medications at the time. Prior to the experiment, the participants were pre-screened for MRI contraindications and had signed a written informed consent form. The study procedure and protocols were approved by the ethics committee of the University of Graz, Austria. The current study represents a part of a bigger project, which was pre-registered prior to data collection (https://aspredicted.org/blind.php?x=SYF_WK6).

### Stimuli and task

Visual stimuli were generated using MATLAB R2019b (MathWorks, Natick, USA) and Psychophysics Toolbox 3 (Brainard, 1997; Kleiner et al., 2007; Pelli, 1997) on a Linux computer (Ubuntu 18.04 LTS). The stimuli were presented in the MR scanner room on a 32” gamma-corrected monitor (resolution: 1920 × 1080 pixels, refresh rate: 60 Hz, maximum surface luminance: 405 cd/m^2^, Nordic NeuroLab, Bergen, Norway), which were viewed via a mirror mounted above the head coil. The total distance from the display to the eyes of the participants was 143.5 cm. Before entering the scanner room, we assessed the participants’ visual acuity with the Snellen chart. We gave the participants a thorough instruction regarding the detection task (see below) and advised them not to move their head during the entire MRI acquisition.

In each experimental block, we presented one of the four possible stimulus conditions (“no illusion”, “diamond”, “left triangle” and “right triangle”, **Figure 1A**) on the screen for a total of 12 seconds. Each stimulus consisted of four black (RGB: 0, 0, 0) pac-man inducer combinations, which were presented against a gray background (RGB: 128, 128, 128). Each inducer consisted of a circle shape with a cut out wedge and subtended 4.084° of visual angle. The inducers were arranged in a diamond shape around the central fixation circle at an eccentricity of 5.774° of visual angle. In the “no illusion” configuration, all inducers were oriented with the 90° cut out wedge facing outwards, thus not creating an illusory shape in the center. In the “diamond” configuration, the inducers were aligned to form a centrally positioned illusory diamond shape. In the “left triangle” configuration, top and bottom inducers had a 45° cut out wedge and were rotated such that an illusory triangle formed within the left visual hemifield. The “right triangle” configuration was a vertically flipped version of the left triangle configuration resulting in an illusory triangle within the right visual hemifield. To account for adaptation in the visual cortex, the stimuli were continuously flickering between pac-man inducers and full circles every 500 ms throughout each experimental block.

**Figure 1.**
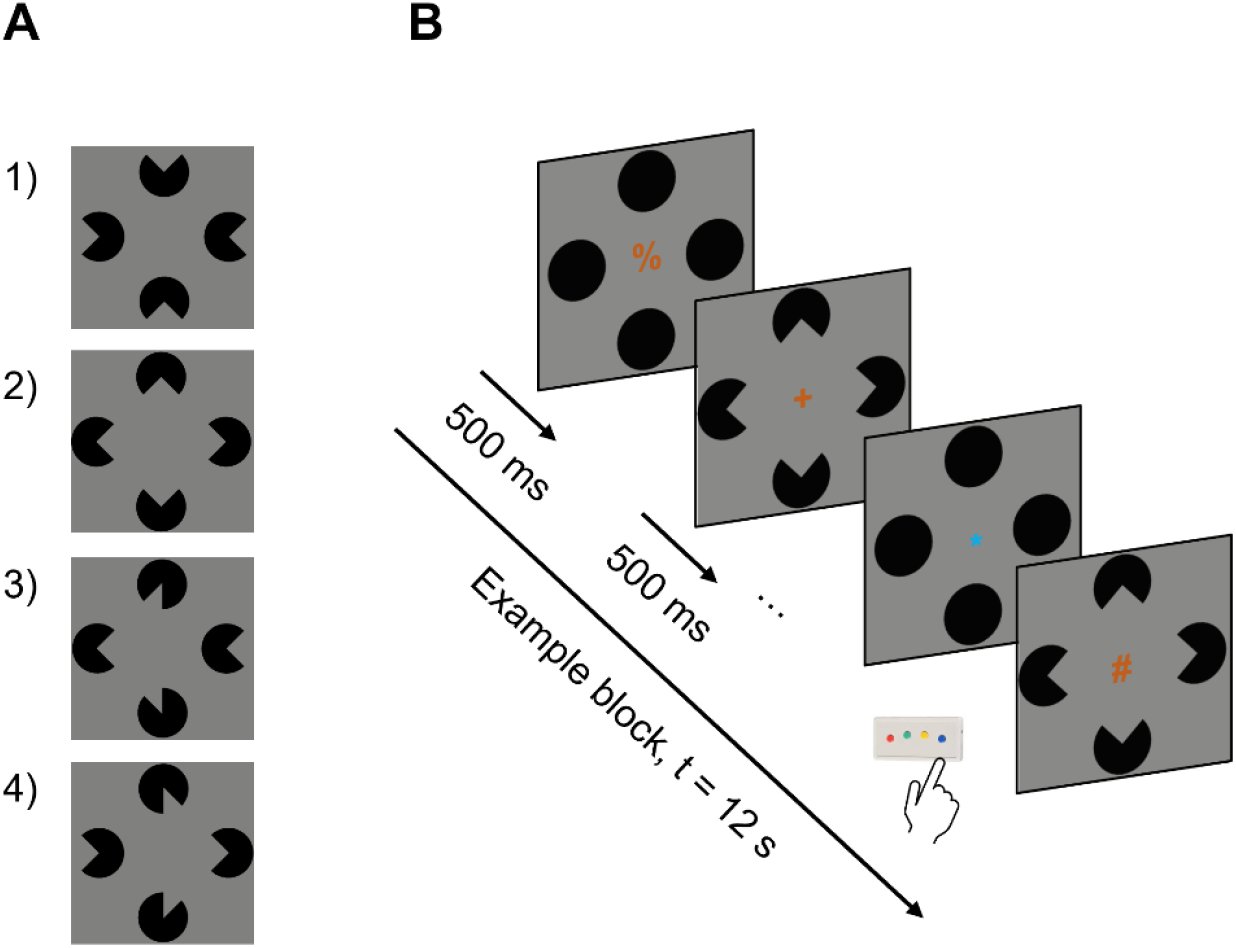
Stimulus types and experimental paradigm. **A**: Stimuli arranged in four possible inducer configurations: 1) no illusion, 2) diamond, 3) left triangle and 4) right triangle. **B**: Sequence of events within one experimental block. The proportions in this figure are altered for illustration purposes. Video examples of blocks for each inducer configuration can be found here:https://osf.io/n9bjr/.

To ensure central gaze fixation and equal distribution of attention across inducer configurations, participants performed a central detection task. A stream of orange (RGB: 153, 80, 0) and blue (RGB: 0, 107, 209) symbols subtending 0.6° of visual angle was presented at the center of the display. A pseudo-randomly chosen symbol was presented one at a time from a fixed set of 26 possible symbols (!”$%&/()=?{[]}\+*∼#-:;><@^). Each symbol was presented for 12 frames (∼ 200 ms) on the screen. The participants were instructed to always attend to the centrally positioned stream of symbols. Participants had to perform one of two tasks. In the first task (“easy”), the participants were instructed to press a button on the response box in their hand each time the symbol color changed to blue (5% of all symbols), regardless of the symbol identity. In the second task (“hard”), the participants were instructed to press the button each time the plus symbol (“+”) appeared on the screen (5% of all symbols), regardless of its color. The symbol stream appearance was thus identical in the two task conditions, and only the task demands varied. The sequence of events within one block can be seen in **Figure 1B**.

Each of the four stimulus conditions (block) was presented six times per run. The sequence of experimental blocks was determined for each run by generating a first-order counterbalanced sequence of conditions (Brooks, 2012). Baseline blocks (12 s each) were added after every fourth experimental block, with a total of six baseline blocks per run. During the baseline blocks, a central fixation point was presented in place of the symbol stream and non-flickering black circles were presented in place of pac-man inducers. In total, there were 30 blocks in each run. The total duration of one run was 360 seconds (excluding the 4 initial dummy scans, which were subsequently discarded). Between each run there was a short (1-2 minutes) break, during which the participants could rest their eyes. In total, one scanning session included eight experimental runs (four runs per task condition). The order of runs with easy and hard detection tasks was pseudorandomized and counterbalanced between the participants. Additionally, after the main experiment we collected an anatomical MRI scan for each participant. The total duration of the whole session was around 60 minutes.

After the MRI acquisition, participants received a questionnaire with six questions for assessment of their subjective visual experience during the experimental paradigm. The participants rated the perceived difficulty of the tasks and the subjective strength of illusory figures on a Likert scale ranging from 1 to 6. The questionnaire included the following questions: “Please rate your subjective perception of the difficulty of the” - Q1: symbol task”; Q2: “color task” (1 – not difficult at all, 6 – very difficult). “During the experiment, how strong did you perceive the following illusory figures?” - Q3: “illusory diamond during the symbol task”; Q4: “illusory diamond during the color task”; Q5: “illusory triangles during the color task”; Q6: illusory triangles during the symbol task” (1 – not strong at all, 6 – very strong).

### MRI data acquisition

Neuroimaging data were acquired using a 3 T Siemens MAGNETOM Vida scanner (Siemens Healthineers, Erlangen, Germany) with a 64-channel head coil. Functional MRI scans were acquired using the simultaneous multi-slice (SMS) accelerated echo-planar imaging (EPI) sequence using T2*-weighted blood oxygenation level-dependent (BOLD) contrast (58 axial slices, TR = 2000 ms, TE = 30 ms, FOV = 220 mm, flip angle = 82°, voxel size = 2.0 × 2.0 × 2.0 mm, SMS factor 2). To compensate for distortion correction during preprocessing, we also acquired one EPI image with opposite phase encoding direction (P >> A). Additionally, a high-resolution atomical scan was acquired using the T1-weighted MPRAGE sequence (TR = 2530 ms, TE = 3.88 ms, TI = 1200 ms, voxel size: 1.0 × 1.0 × 1.0 mm, GRAPPA factor 2).

### MRI data preprocessing

Before any data preprocessing, we removed facial identifiers from all participants with pydeface 2.0.0 (Gulban et al., 2019). Afterwards, we visually inspected the quality of MRI images with MRIQC 0.16.1 (Esteban et al., 2017). Anatomical and functional MRI data were preprocessed using fMRIPrep 20.2.3 (Esteban et al., 2019), which is based on Nipype 1.6.1 (Gorgolewski et al., 2011). The main preprocessing steps are summarized below. Detailed notes about the preprocessing pipeline are available here: https://osf.io/n9bjr/.

#### Anatomical data preprocessing

The T1-weighted (T1w) image was corrected for intensity non-uniformity with ANTs *N4BiasFieldCorrection* function and used as T1w-reference, which was then skull-stripped. Cerebrospinal fluid (CSF), white-matter (WM) and grey-matter (GM) brain tissue segmentation was performed on the brain-extracted T1w using FSL *fast* function from FSL 5.0.9 (Woolrich et al., 2009). Brain surfaces were reconstructed using FreeSurfer *recon-all* function (Dale et al., 1999).

#### Functional data preprocessing

First, a reference volume and its skull-stripped version were generated using a custom methodology of fMRIPrep. For each BOLD run the following preprocessing transformations were performed: A B0-nonuniformity map was estimated based on the EPI with opposite phase-encoding direction using *3dQwarp* from AFNI 20160207 (Cox, 1996). Based on the estimated susceptibility distortion, a corrected EPI reference was calculated for more accurate co-registration with the anatomical reference. The BOLD reference image was co-registered to the T1w reference image using FreeSurfer’s *bbregister* (Greve & Fischl, 2009). Head-motion parameters with respect to the BOLD reference image were estimated before any spatiotemporal filtering using *mcflirt* (FSL 5.0.9). Slice-time correction was performed using *3dTshift* from AFNI 20160207. These transforms were concatenated and applied in one interpolation step. The BOLD-time series were resampled to the fsaverage surface using *mri_vol2surf* (FreeSurfer). For the region-of-interest (ROI) general linear model (GLM) analyses (see below), we used unsmoothed preprocessed BOLD runs. For the whole-brain GLM analyses, each preprocessed BOLD run was smoothed along the cortical surface with a 2D Gaussian kernel of 5 mm full-width at half-maximum (FWHM) using *mri_surf2surf* (FreeSurfer).

### Data analysis

#### First-level GLM analysis

All first-level GLM analyses were performed using FsFast (FreeSurfer Functional Analysis Stream). For each participant, we performed a standard GLM analysis, with each experimental condition modelled as a unique regressor convolved with a canonical hemodynamic response function (HRF). In contrast to early visual areas, areas of the superior parietal cortex are known to show a transient BOLD response pattern (Steinman et al., 1997; Yantis et al., 2002; Zaretskaya et al., 2013), which we confirmed for our paradigm with an additional FIR GLM analysis. Thus, we modelled the response function into the GLM as 0.1 s events at block onsets rather than full 12 s block durations. There were eight unique condition regressors in total, with the first four modelling stimulus conditions under the easy task and the next four under the hard task. Additionally, run-specific effects as well as slow signal drifts were added as nuisance regressors. Illusory shape activity was quantified by calculating six contrast estimates (CES). The latter were defined as the difference of β-estimates between each illusory shape condition (diamond, left triangle and right triangle) and the “no illusion” condition, separately for each task type. The same GLM was fitted to the unsmoothed data for the ROI analysis and to the smoothed data for an additional whole-brain group analysis. All statistical analyses for ROI and behavioral data were performed in RStudio 4.1.2 (RStudio Team, 2021).

#### Definition of regions of interest and ROI analysis

Seven topographic areas of the posterior parietal cortex (IPS0, IPS1, IPS2, IPS3, IPS4, IPS5 and SPL1) were defined using the maximum probability maps (MPM) of the surface-based probabilistic atlas by Wang et al. (2015), resampled to FreeSurfer fsaverage standard space (Benson & Winawer, 2018). Since our study is hypothesis-driven, an a priori definition of the parietal areas via probabilistic MPMs is a recommended method for inferring about the individual’s anatomical locations within the brain (Eickhoff et al., 2006). Furthermore, the predefined areas in a standard surface space allow for unbiased and direct comparison with data from other studies (Wang et al., 2015). **Figure 2A** illustrates their location on an inflated cortical surface. Number of vertices for each parietal ROI were: IPS0 = 1055, IPS1 = 900, IPS2 = 899, IPS3 = 604, IPS4 = 148, IPS5 = 53, and SPL1 = 232 vertices in the left hemisphere and IPS0 = 835, IPS1 = 675, IPS2 = 624, IPS3 = 805, IPS4 = 160, IPS5 = 18, and SPL1 = 295 vertices in the right hemisphere. We extracted mean vertex values within each ROI for each contrast estimate map of each participant. The per-ROI means were then used to conduct statistical analysis. Since we do not expect a lateralization of the illusory diamond response, we averaged mean shape response over both hemispheres. For the illusory triangle response, we averaged the two contralateral triangle values (i.e., the left hemisphere value for the contrast “right triangle vs. no illusion” and the right hemisphere value for the contrast “left triangle > no illusion”) and the two ipsilateral values (i.e., the left hemisphere value for the contrast “left triangle vs. no illusion” and the right hemisphere value for the contrast “right triangle > no illusion”).

**Figure 2.**
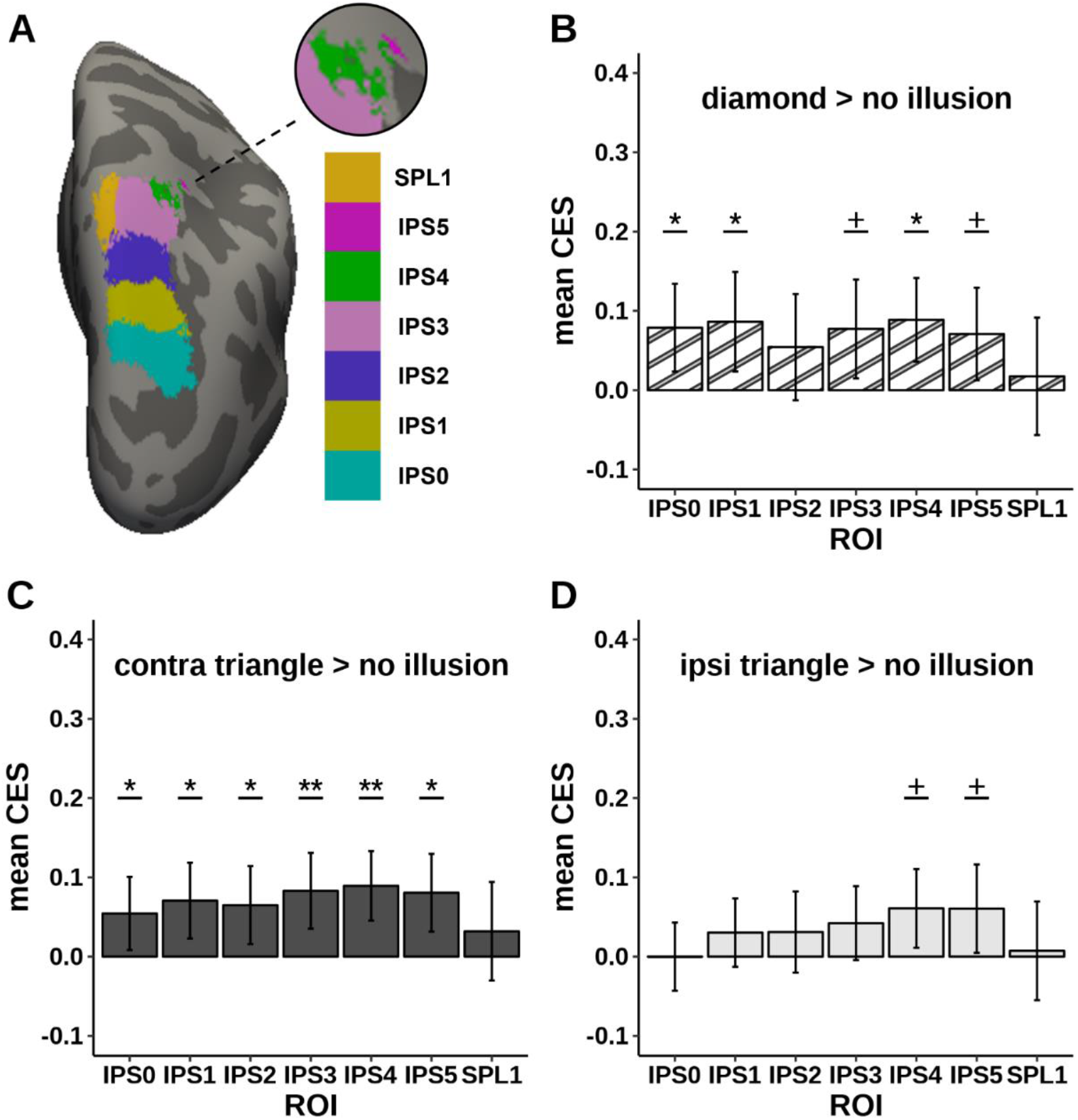
**A:** Location of the parietal areas from the probabilistic surface-based atlas (Wang et al., 2015), overlayed onto the right hemisphere of the inflated fsaverage cortical surface. Inset shows enlarged anterior areas IPS4 and IPS5. **B-D:** Illusory shape-related response during the easy task within the posterior parietal cortex (B: diamond, both hemispheres; C: triangle, contralateral hemisphere; D: triangle, ipsilateral hemisphere). Error bars represent 95% CIs. ^+^*p*<.05, uncorrected; \**p*<.05, \*\**p*<.01, Holm-Bonferroni corrected (7 tests).

The following statistical tests were carried out to investigate the illusory shape response in the parietal ROIs. First, we tested the presence of illusory shape responses in each parietal ROI by testing for non-zero contrast differences (“diamond > no illusion”, “contralateral triangle > no illusion” and “ipsilateral triangle > no illusion” during the easy task) with a two-tailed one sample t-test. Second, we tested the spatial specificity of the illusory shape response within each ROI by comparing the contralateral and the ipsilateral responses during the easy task with a two-tailed paired samples t-test. Third, we tested whether higher task demands affected the illusory diamond response by directly comparing the illusory diamond response for the two task difficulties with a two-tailed paired samples t-test within each ROI. Finally, we tested whether higher task demands affected illusory triangle response by performing a 2 × 2 repeated-measures ANOVA within each ROI, with laterality (contralateral and ipsilateral) and task (easy and hard) as factors.

#### Additional whole-brain group analysis

To confirm previous findings on illusory shape response in ventral and early visual areas we performed an additional whole-brain group analysis. The analysis was performed using FreeSurfer *mri_glmfit* command with -wls flag (weighted-least-squares), testing for a nonzero difference between each illusory shape and no illusion conditions, separately for each task. Correction for multiple comparisons was performed using precomputed z-based Monte Carlo simulation with a cluster-wise threshold at p < .05 (Hagler et al., 2006) and a cluster-forming threshold at p < .001 to minimize false positive rates (see Greve & Fischl, 2018). Finally, we adjusted p-values with a Bonferroni correction for two spaces (left and right hemisphere). The surviving clusters were classified according to the Desikan-Killiany cortical parcellation (Desikan et al., 2006). Additionally, we calculated the overlap between the clusters and areas of the visual probabilistic atlas (Wang et al., 2015).

#### Behavioral data analysis

In our behavioral data analysis, we ensured that our manipulation of the task difficulty was successful. The difficulty of both detection tasks was verified with the average hit rate and the participants’ subjective rating of the task difficulty. The hit rate in each run was calculated as a total number of correctly detected target instances (target color or target symbol, depending on the task), divided by the total number of target instances and multiplied by 100. Values from the runs with the same task type were averaged.

We compared the average hit rates between the easy and hard task and the subjective ratings of the task difficulty by performing a two-tailed paired samples Wilcoxon signed rank test with continuity correction. To determine whether differences in illusory shape response between tasks observed in fMRI can be explained by differences in subjective experience of the illusory shape, we also compared participants’ ratings of their subjective illusion perception between tasks with two Wilcoxon signed rank tests, one for the diamond and one for the triangle shape.

#### Data and code availability

ROI and group-level fMRI, behavioral results, as well as control analyses and relevant scripts are available at the OSF project repository:https://osf.io/n9bjr/. Our institution’s data protection policy does not allow publicly sharing native space MRI data.

## Results

### Illusory shape response

We first examined whether parietal topographic regions responded to illusory Kanizsa shapes in the easy task, which is comparable to a typical fixation task used in fMRI. We performed a one-sample t-test for a non-zero illusory diamond response (difference between the illusory diamond and no illusion) in each ROI. Significant illusory shape response was present in areas IPS0, 1, 3, 4 and 5. SPL1 and IPS2 did not respond to the illusory diamond (**Figure *2*B**).

The surface of the illusory diamond was occupying the central visual field around the fixation task. It was therefore easy to perceive and potentially captured attention. We therefore also tested if illusory shape responses were present in the parietal areas for the triangles which were placed peripherally relative to the central task. All six ROIs of the intraparietal sulcus (IPS0-5) responded to the contralateral illusory triangle during the easy task. SPL1 was not activated by the illusory triangle (**Figure *2*C**). Interestingly, IPS4 and IPS5 also responded to the illusory triangle presented to the ipsilateral hemisphere (**Figure 2D**). Statistical details about illusory shape responses are presented in **Table 1**.

**Table 1.**
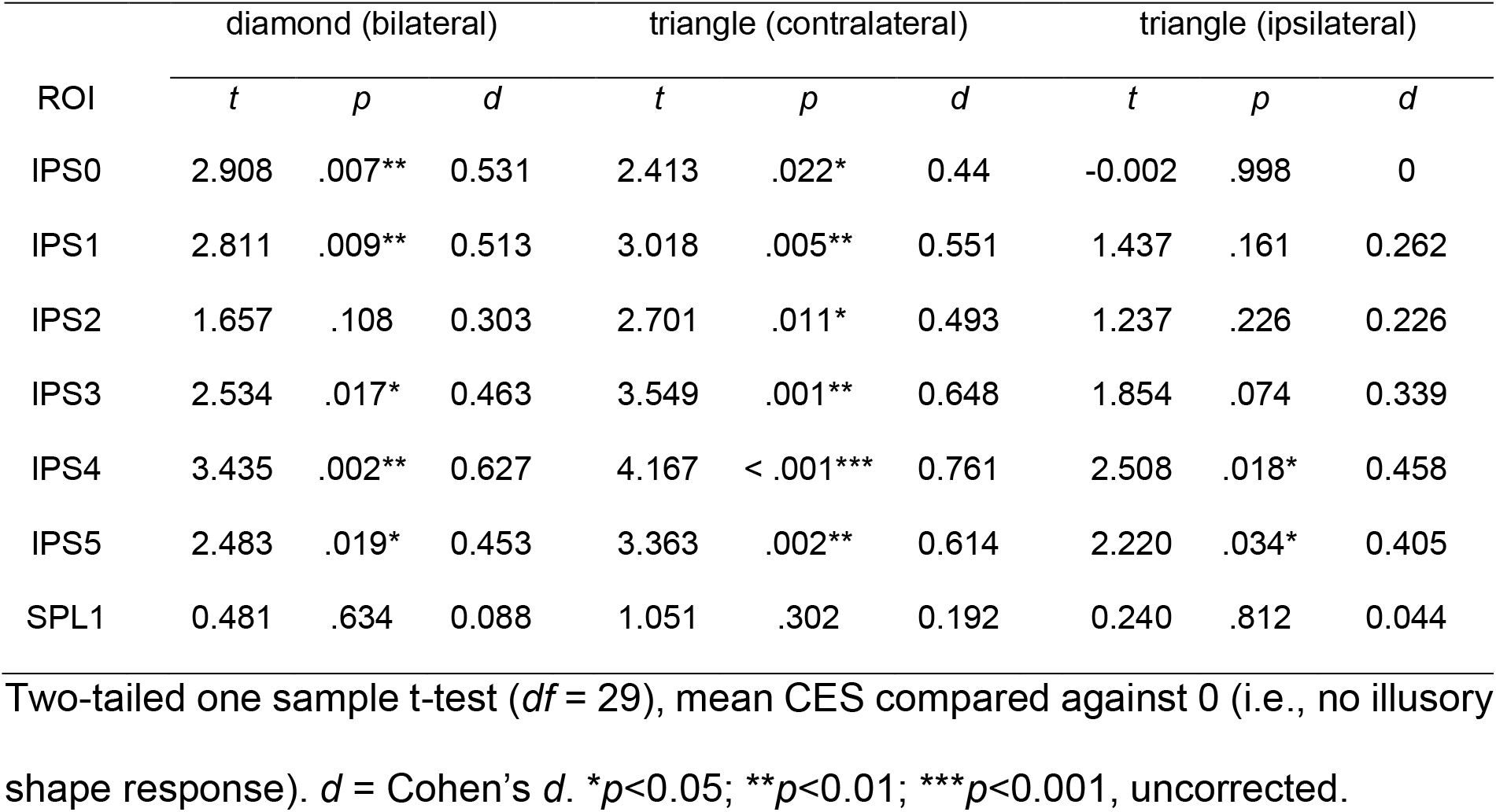
Illusory shape response (shape > no illusion, easy task).

| ROI | diamond (bilateral) |  |  | triangle (contralateral) |  |  | triangle (ipsilateral) |  |  |
| --- | --- | --- | --- | --- | --- | --- | --- | --- | --- |
|  | *t* | *p* | *d* | *t* | *p* | *d* | *t* | *p* | *d* |
| IPS0 | 2.908 | .007** | 0.531 | 2.413 | .022* | 0.44 | -0.002 | .998 | 0 |
| IPS1 | 2.811 | .009** | 0.513 | 3.018 | .005** | 0.551 | 1.437 | .161 | 0.262 |
| IPS2 | 1.657 | .108 | 0.303 | 2.701 | .011* | 0.493 | 1.237 | .226 | 0.226 |
| IPS3 | 2.534 | .017* | 0.463 | 3.549 | .001** | 0.648 | 1.854 | .074 | 0.339 |
| IPS4 | 3.435 | .002** | 0.627 | 4.167 | < .001*** | 0.761 | 2.508 | .018* | 0.458 |
| IPS5 | 2.483 | .019* | 0.453 | 3.363 | .002** | 0.614 | 2.220 | .034* | 0.405 |
| SPL1 | 0.481 | .634 | 0.088 | 1.051 | .302 | 0.192 | 0.240 | .812 | 0.044 |

Two-tailed one sample t-test ( $df = 29$ ), mean CES compared against 0 (i.e., no illusory shape response). $d$ = Cohen's $d$ . $*p < 0.05$ ; $**p < 0.01$ ; $***p < 0.001$ , uncorrected.

### Spatial specificity of the illusory shape response

Next, we tested whether illusory shape responses were spatially specific by directly comparing responses to the contralateral with responses to the ipsilateral triangles using two-tailed paired samples t-tests for each ROI. Contralateral responses were stronger in IPS0 (*t*(29) = 6.397, *p* < .001, *d* = 1.17), IPS1 (*t*(29) = 4.090, *p* < .001, *d* = 0.747), IPS2 (*t*(29) = 3.203, *p* = .003, *d* = 0.585), IPS3 (*t*(29) = 3.461, *p* = .002, *d* = 0.632) and SPL1 (*t*(29) = 2.109, *p* = .044, *d* = 0.385). We found no hemispheric difference in illusory triangle shape response in IPS4 (*t*(29) = 1.724, *p* = .095, *d* = 0.315) or IPS5 (*t*(29) = 0.964, *p* = .343, *d* = 0.176). The comparison between contralateral and ipsilateral responses is shown in **Figure 3**.

**Figure 3.**
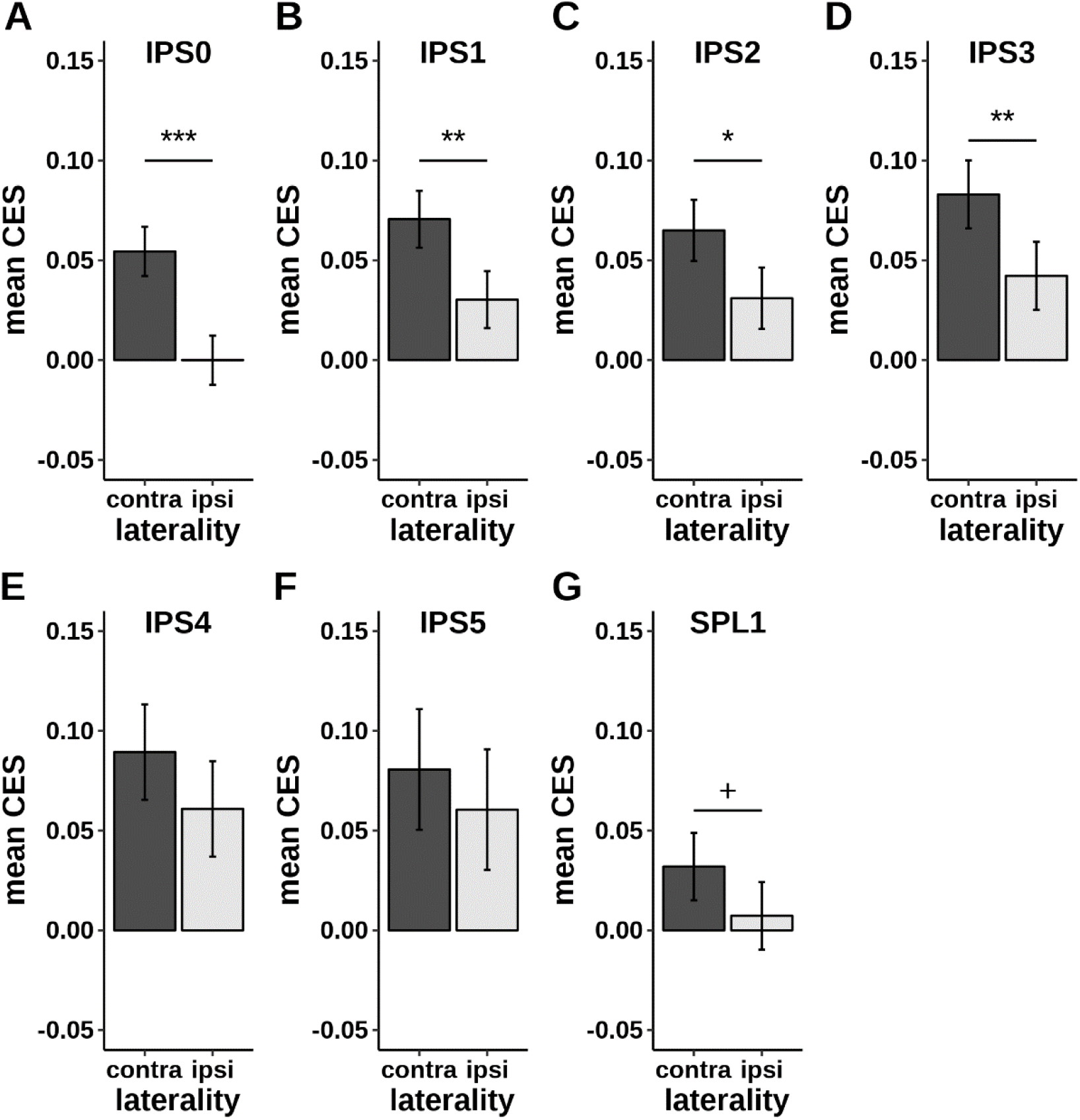
**A-G:** Spatial specificity of illusory triangle response (easy task). Error bars represent 95% CIs (Morey, 2008). ^+^*p*<.05, uncorrected; \**p*<.05, \*\**p*<.01, \*\*\**p*<.001, Holm-Bonferroni corrected (7 tests).

### Attentional modulation of the illusory shape response

Finally, we tested whether the observed illusory shape response was modulated by task difficulty by comparing the responses in the easy task with responses in the hard task. Our 2×2 repeated measures ANOVA revealed that shape-related response in IPS0, IPS1, IPS2, and IPS3 showed preference to the contralateral stimulus presentation (main effect of laterality IPS0: *F*(1, 29) = 48.628, *p* < .001, *η*_*p*_^*2*^ = .626; IPS1: *F*(1, 29) = 21.705, *p* < .001, *η*_*p*_^*2*^ = .428; IPS2: *F*(1, 29) = 10.908, *p* = .003, *η*_*p*_^*2*^ = .273; IPS3: *F*(1, 29) = 14.632, *p* < .001, *η*_*p*_^*2*^ = .335, see **Figure 4A-D**). Crucially, the illusory triangle responses were not attenuated by the hard task and there was no interaction between task and laterality. In contrast, IPS4 showed an attenuation of the illusory triangle response during the hard task compared to the easy task (main effect of the task *F*(1, 29) = 7.726, *p* = .009, *η*_*p*_^*2*^ = .21, **Figure 4E**). Crucially, this attenuation occurred irrespective of laterality (no main effect of laterality) and there was no interaction between task and laterality. IPS5 revealed a trend in the same direction as IPS4 (**Figure 4F**), but it did not reach significance (*F*(1, 29) = 3.795, *p* = .061, *η*_*p*_^*2*^ = .116). SPL1 did not show any significant effects. **Figure 4G** shows the visual representation of location of the main effects for each IPS area.

**Figure 4.**
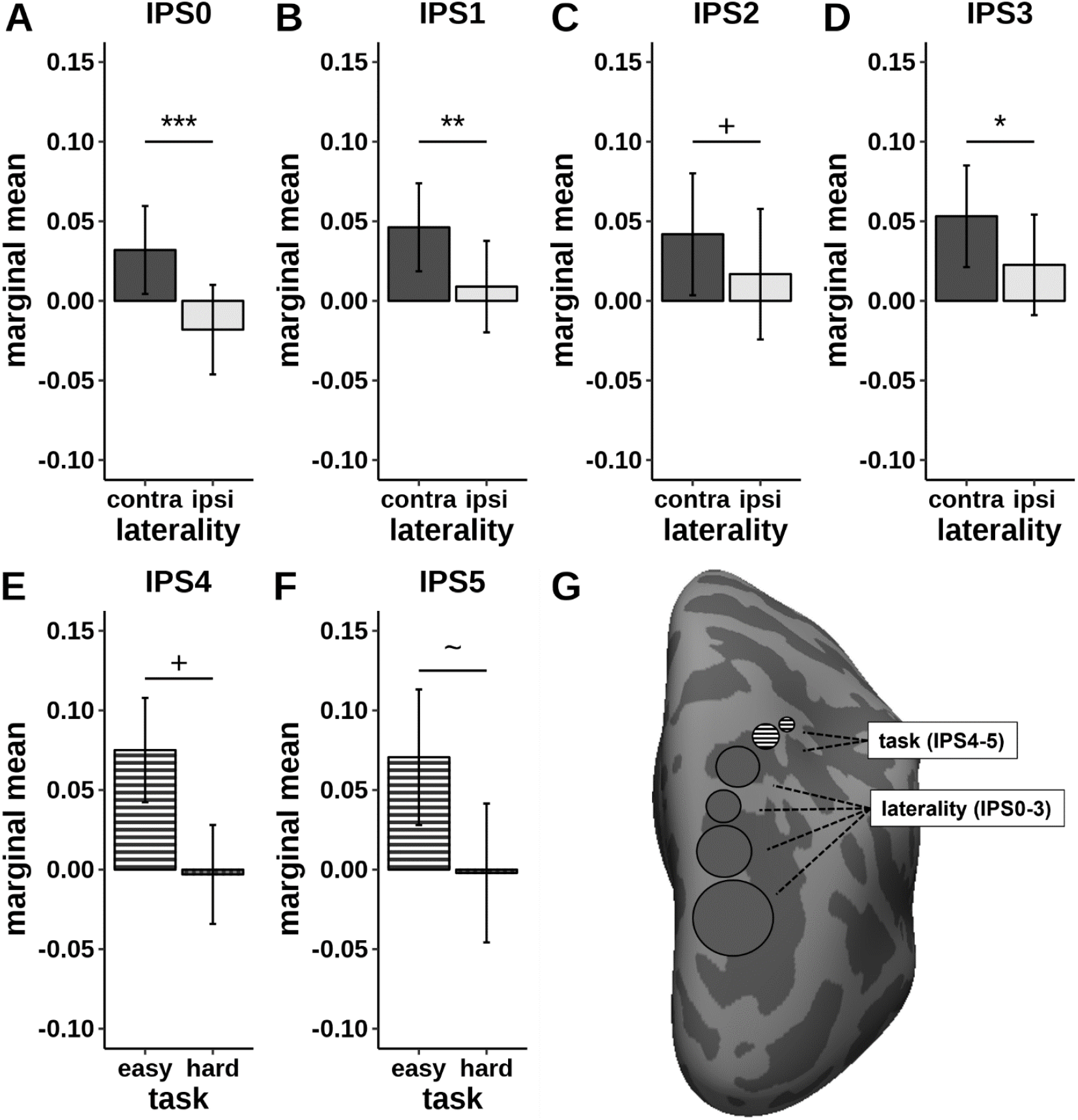
Region-of-interest parietal brain responses and visualization of their respective cortical surface location. **A-D:** Main effect of laterality in areas IPS0-3. **E-F:** Main effect of task in area IPS4 and trend of main effect of task in IPS5. Error bars represent 95% CIs (Morey, 2008). ∼*p*<0.1, +p<.05, uncorrected. \**p*<.05, \*\**p*<.01, \*\*\**p*<.001, Holm-Bonferroni corrected (21 effects). **G:** Visualization of main effects of laterality and task within the posterior parietal cortex on the right inflated cortical surface. Circle size is proportional to respective effect size.

To determine whether higher attentional demands reduced the illusory diamond response, we compared the responses across different task difficulties with a two-tailed paired-samples t-test. However, we found no effect of task on the illusory diamond response in any of the seven parietal ROIs.

### Behavioral results

Our behavioral results confirmed that our manipulation of task difficulty was successful. We found a significant difference in hit rates between the two detection tasks (*Z* = 4.772, *p* < .001, *r* = 0.873), with the easy task runs having higher hit rates than the hard task runs (**Figure 5A**). Corresponding effects were found for the subjective difficulty ratings as well, with the easy task being perceived as easier compared to the hard task (*Z* = -3.818, *p* < .001, *r* = 0.716, **Figure 5B**). Another two separate Wilcoxon signed ranked tests with continuity correction comparing subjective illusion strength revealed no difference between the tasks (diamond: *Z* = 0.133, *p* = .894, *r* = 0.076; triangle: *Z* = -0.619, *p* = .536, *r* = 0.111).

**Figure 5.**
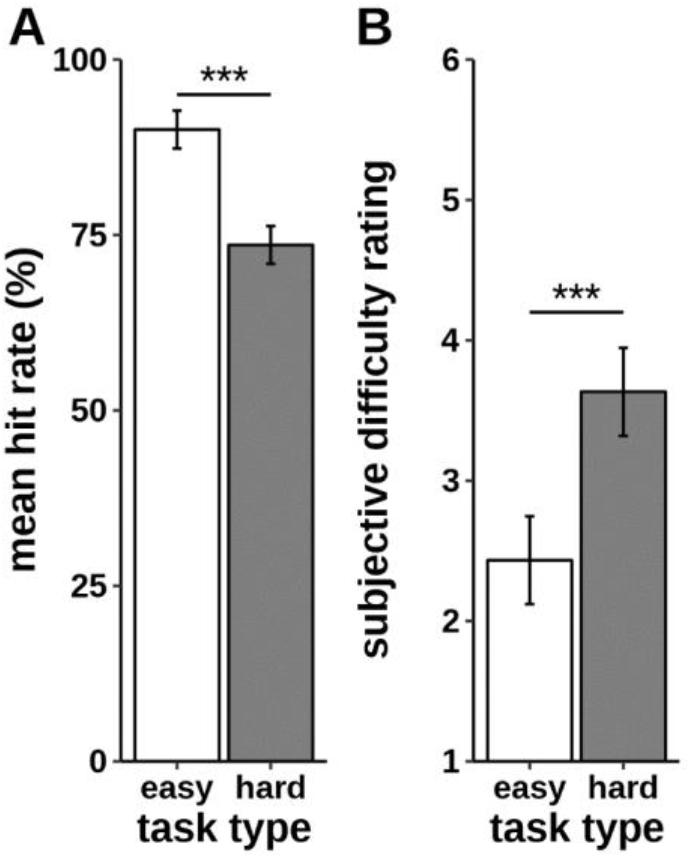
Objective and subjective measurements of task difficulty. **A**: Mean hit rates in easy and hard detection tasks. **B:** Mean ratings of subjective task difficulty, rated on a 6-point Likert scale (1 – not difficult at all, 6 – very difficult). Error bars represent 95% CIs (Morey, 2008). ***p<.001.

### Whole-brain results

Whole-brain group results are shown in **Figure 6**. We found shape-related responses in early visual cortex (V1-V3), in areas hV4, V3b and LO1 across all illusory conditions, confirming findings of multiple previous studies. The increase of task difficulty reduced activation and increased deactivation in all illusory conditions. All significant clusters and their respective sizes and coordinates can be found here:https://osf.io/n9bjr/. Positive cluster overlaps with the areas of the probabilistic visual atlas can be seen in **Table 2**. Only one of the clusters that survived correction for multiple comparisons was located in the parietal cortex (illusory diamond in the easy task, cluster size: 243 vertices, peak MNI coordinates: [22.8, -56.8, 55.8]). The cluster overlaps with the surface area of IPS3 (182 vertices, 74.9% of total ROI size) and IPS4 (57 vertices, 23.46% of total ROI size). For the sake of comparison with previous findings, we also compared the location of this parietal cluster with two parietal clusters related to illusory shape perception reported in Zaretskaya et al. (2013). 177 vertices of the “SPL” cluster from their study (14.41% of the total “SPL” size) overlaps with the parietal cluster, and there is no overlap with the “aIPS” cluster from the same study.

**Table 2.** Percentage overlap of Figure 6 shape-activated clusters (positive clusters only) with visual topographic areas.

|  | diamond |  |  |  | contralateral triangle |  |  |  |
| --- | --- | --- | --- | --- | --- | --- | --- | --- |
|  | easy LH | hard LH | easy RH | hard RH | easy LH | hard LH | easy RH | hard RH |
| V1 | 23.048 | 18.634 | 27.389 | 19.876 | 20.539 | 15.520 | 17.404 | 13.457 |
| V2 | 27.461 | 25.213 | 31.208 | 27.201 | 30.377 | 23.572 | 29.740 | 23.138 |
| V3 | 42.355 | 35.780 | 31.615 | 24.385 | 41.667 | 34.709 | 34.846 | 24.154 |
| hV4 | 41.723 | 20.635 | 49.409 | 21.260 | 33.333 | 21.995 | 41.929 | 14.370 |
| V3a | 3.852 | - | - | - | 4.708 | 2.710 | - | - |
| V3b | 77.828 | 50.679 | 61.354 | 24.017 | 48.190 | 12.670 | 12.445 | 5.895 |
| LO1 | 94.457 | 23.725 | 93.345 | 46.585 | 56.319 | 31.929 | 60.946 | 28.371 |
| LO2 | 20.110 | - | 28.105 | - | - | - | - | - |
| IPS3 | - | - | 22.609 | - | - | - | - | - |
| IPS4 | - | - | 35.625 | - | - | - | - | - |

Reported vertices in V1-V3 represent combined dorsal and ventral part of the respective ROI. Only cluster overlaps with $\geq 10$ vertices within a ROI are shown. Values represent vertex overlap as a percentage of the respective overlapping ROI size: $(\text{cluster}_n / \text{ROI}_n) * 100$ .

**Figure 6.**
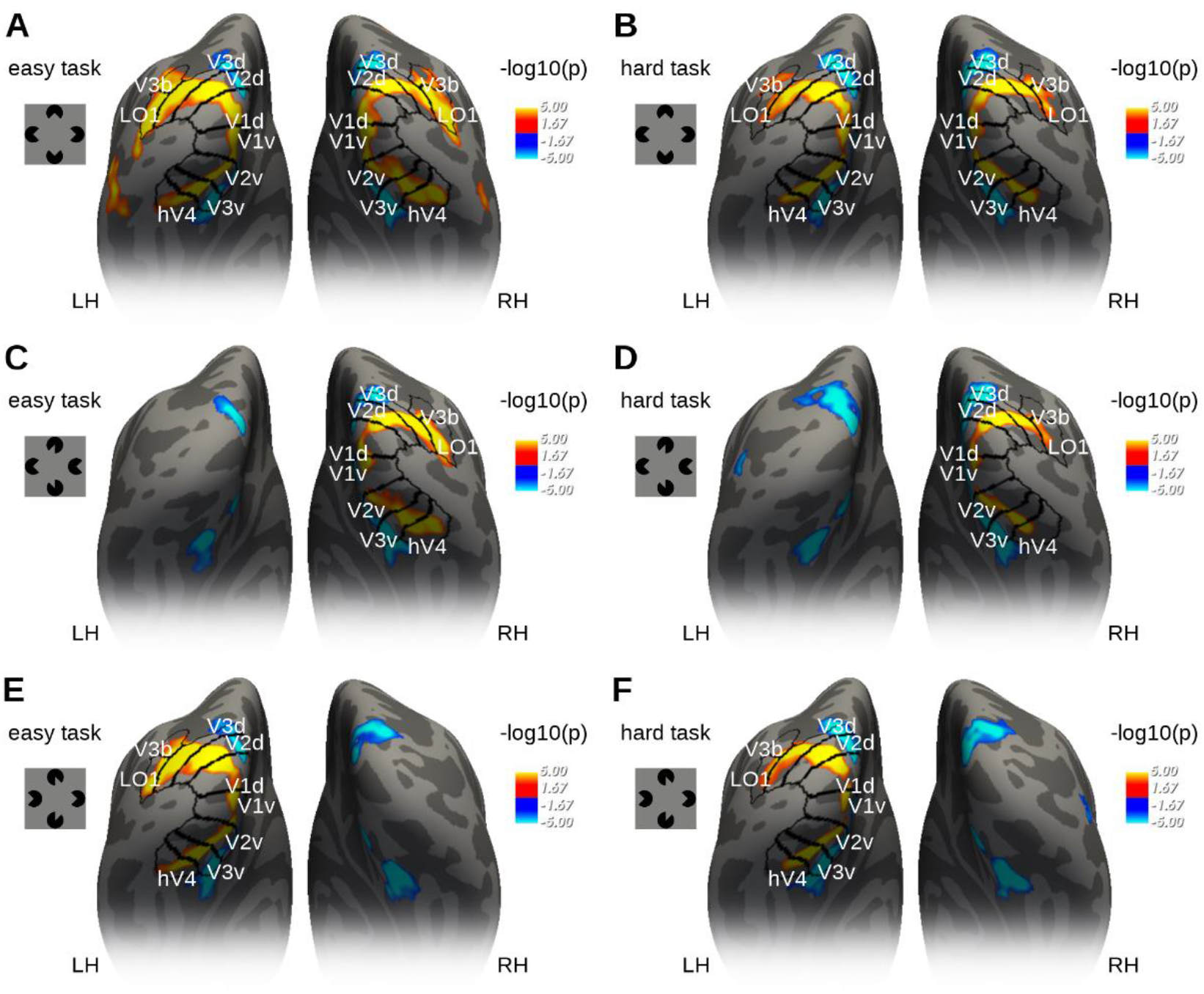
Whole-brain group-level results for each illusory shape > no illusion contrasts (corrected for multiple comparisons). **A**: diamond, easy task; **B**: diamond, hard task; **C**: left triangle, easy task; **D**: left triangle, hard task; **E**: right triangle, easy task; **F**: right triangle, hard task. Clusters of significant responses are shown on the inflated cortical surface of the fsaverage template. LH – left hemisphere. RH – right hemisphere.

## Discussion

In this study we examined responses of parietal topographic areas to illusory Kanizsa shapes. Our results show that IPS areas respond to illusory shapes even when they are not defined by motion. The results also indicate that different parietal areas respond to illusory shapes in a different manner. The posterior areas IPS0 to IPS3 show a spatially specific illusory shape response, which is not present in the anterior IPS areas. On the other hand, the anterior areas IPS4 and IPS5 show an attention-dependent illusory shape response, which is not the case for the posterior areas. Our results thus also show a dissociation between the topographically specific and attention-dependent activation during illusory shape perception along the posterior-anterior gradient within the IPS.

### Static illusory shape responses in the parietal cortex

Several previous fMRI studies already reported illusory shape response within the superior parietal cortex and the intraparietal sulcus, but those studies relied on motion-defined bistable stimuli which either did or did not produce an impression of a moving illusory shape (Grassi et al., 2016, 2018; Zaretskaya et al., 2013). A bistable paradigm has the advantage of fully matching the illusory and non-illusory conditions. However, it requires participants to track their own subjective experience of the stimulus, so it inevitably leads to a coupling between illusory perception and attention. Due to this coupling, previous studies could test neither the topographic specificity of the shape responses, nor their dependence on attention. In the current study, highly controlled Kanizsa stimuli allowed us to overcome these constraints, decoupling the spatial location of an illusory shape from the locus of attention. We demonstrate that illusory content of the Kanizsa shapes is represented within the posterior maps IPS0-3 even when attention is controlled for. This finding implies that parietal topographic areas play a general role in representing illusory objects irrespective of the exact perceptual mechanism, be it motion or inducer alignment, that led to their generation.

### Spatial specificity of shape responses

The decupling of illusory shape location from the locus of attention revealed that the response of the most posterior IPS regions (IPS0-3) is spatially specific. These areas respond stronger to the illusory triangles presented to the contralateral compared to ipsilateral side of the visual field. Since each IPS map represents the contralateral visual field, such specificity implies that illusory content in these areas is represented, at least on a coarse spatial scale, in a topographic manner. The topography of the illusory shape response and its independence from attention in these areas resembles the responses of early visual areas V1 and V2, which show a differentiated pattern of illusory surface enhancement and inducer/background suppression (Grassi et al., 2017; Kok & de Lange, 2014, but see de-Wit et al., 2012). Due to the relatively small surface area and much larger receptive field sizes of the IPS maps compared to V1/V2, it is challenging to show such fine-scale response differences with conventional fMRI. Future studies utilizing high-resolution fMRI at ultrahigh field have the potential to reliably distinguish between the different sub-parts of the Kanizsa figure within the parietal topographic maps and test whether there is a distinction between illusory surface enhancement and inducer suppression (Zaretskaya, 2021). Additionally, such studies could also focus on the distribution of illusory shape responses across the cortical depth to determine the feed-forward vs. feedback origin of these signals (Kok et al., 2016).

Interestingly, spatial specificity was not found in the most anterior maps IPS4 and IPS5. The most likely explanation is the increasing receptive field sizes of neurons along the visual hierarchy, which is also present within the IPS regions (Klein et al., 2014). Large receptive field sizes that span both hemifields within the anterior IPS maps could have led to nearly equal ipsi- and contralateral illusory shape responses, providing no net difference between the sides. This explanation is supported by a previous study investigating the effect of attention on population receptive field properties within parietal topographic maps (Sheremata & Silver, 2015). Regardless of whether attention was on the stimulus or not, the authors found reduced contralateral bias of the visual field representation in the anterior IPS regions.

### Dissociation of attention- and illusion-related response

Another major finding of our study is that the attentional modulation of illusory shape responses was confined to the anterior map IPS4, with a trend in IPS5. These two most anterior IPS maps showed a complete absence of responses to peripheral illusory triangles during the more difficult task which forced the allocation of attentional resources to the fovea. Responses of the anterior IPS areas may thus reflect something beyond a mere illusory shape presence. For example, increased attentional demands at the fovea could have led to an inattention blindness-like effect, affecting participants’ ability to consciously perceive the illusory shape (Vandenbroucke et al., 2014). If so, the two anterior IPS regions could represent the conscious, subjective aspects of the Kanizsa illusion, while the remaining areas could signal the automatic preattentive shape response (Conci et al., 2009; Moore & Egeth, 1997; Vuilleumier et al., 2001). Our experimental paradigm was designed to dissociate attention and spatial location of illusory shapes, which prevented us from systematically assessing the subjective illusion strength experienced under each task condition. Although participants reported their subjective experience after the experiment, we did not find any differences in their reports. This question could be addressed in future studies with experimental paradigms that are designed to dissociate awareness- and attention-related processing (Cohen et al., 2020; Pitts et al., 2012, 2018).

Contrary to the illusory triangles, we found no attentional modulation for the illusory diamond. This is expected, since in the case of illusory diamond the locus of attention and the location of the illusory figure coincide. Attentional resources dedicated to the central task could thus also be used to process the foveal illusory diamond surface (Bakar et al., 2008).

### Illusory shape response and predictive coding

Illusory shape responses are frequently interpreted in the context of the predictive coding theory (Friston, 2018; Kok & de Lange, 2015; Rao & Ballard, 1999). According to this theory, the response in the topographic representation of the illusory surface in early visual areas increases because the feedback signal contains predictions that do not match with the bottom-up sensory input, resulting in a prediction error. At the same time, the response in the topographic representation of the inducers decreases because there is a match between the prediction and the sensory input (Grassi et al., 2017; Kok & de Lange, 2014). Our findings in the IPS suggest that the topographic illusory shape-related response is present in multiple areas far beyond EVC. This is consistent with the idea that predictive inference is hierarchical, and that the prediction and prediction error signals propagate through multiple levels of the visual hierarchy (Kok & de Lange, 2015; Muckli & Petro, 2013). The IPS could be enhancing the illusory surface representation on a coarser level and sending back shape information to early visual areas, which are capable of finer and more detailed representation of the subjective content (Roelfsema & de Lange, 2016).

Although the top-down feedback signals are assumed to explain the EVC response profile, it is not known in which areas such feedback signals could originate. One candidate region for the feedback source is the LOC (Chen et al., 2021), which is well known to respond to real (Kourtzi & Kanwisher, 2001) and illusory shapes (Chen et al., 2020; Fang et al., 2008). Our whole-brain results confirm the presence of illusory shape response in both early visual areas and the LOC (LO1). Importantly, our results point to a potential third player in this context, the intraparietal sulcus. Previous bistable perception studies already hypothesized that the IPS could be an alternative candidate for sending the feedback signal to V1, especially because it was active even for stimuli that did not evoke the LOC response (Grassi et al., 2018). Interestingly, studies using bistable stimuli also found a dissociation between the roles of the more anterior and the more posterior IPS regions in bistable perception, with the more anterior IPS carrying the prediction signals (Kanai et al., 2011; Zaretskaya et al., 2013), and the more posterior IPS conveying the prediction error (Kanai et al., 2011). Our results do not support the notion that anterior IPS maps provide the source for the prediction signal, which should have been present in our experiment irrespective of attentional load. Nevertheless, they highlight the functional dissociation within the IPS maps along the posterior-anterior gradient, which is worth further investigation.

## Conclusion

Our study shows that the topographic areas along the intraparietal sulcus represent illusory content, with response in the posterior IPS being spatially specific and the response in the anterior IPS being dependent on attentional resources. The observed illusory shape responses may serve as a feedback signal for the areas earlier in the visual hierarchy. It remains to be seen whether and how responses within the IPS interact with the well-documented shape responses found in the ventral and early visual areas.

## Acknowledgements

This work was supported by the BioTechMed-Graz Young researcher group grant. The authors would like to thank Thomas Zussner for his help during MR data acquisition.

